# Biomimetic scaffolds mimic human subretinal debris for preclinical testing of prosthetic vision

**DOI:** 10.64898/2026.10.06.756601

**Authors:** Mohajeet B Bhuckory, Vladimir Mamchik, Keith Ly, Nathan Jensen, Viktoryia Shautsova, Quentin Devaud, Andrew Shin, Davis Pham-Howard, Roopa Dalal, Ludwig Galambos, Theodore Kamins, Daniel Palanker

## Abstract

Photovoltaic subretinal prostheses replace the lost photoreceptors in patients blinded by atrophic age-related macular degeneration (AMD) and provide central vision by electrically stimulating the inner retinal neurons. Subretinal debris in geographic atrophy separate the inner nuclear layer from the implant by tens of micrometers, increasing the stimulation threshold and reducing the pattern contrast. Absence of this separating layer in animal models diminishes the accuracy of preclinical testing of the next-generation implants, especially with smaller pixels. We developed three-dimensional biomimetic scaffolds nanoprinted onto photovoltaic arrays to provide a precisely controlled separation between the implants and target neurons in animal models of retinal degeneration. Gap sizes of the scaffolds were adjusted to meet the following requirements: they should allow light penetration to the implant, provide unobstructed flow of electric current, but preclude migration of cells into their 3-D structure. Gaps of 3 µm were found to be optimal as they restricted cellular migration and maintained the inner nuclear layer above both planar and pillar electrode arrays. Scaffolds did not measurably alter the photovoltaic response and accurately reproduced the distance dependence of stimulation threshold. Following implantation, the initially fluid-filled intra-scaffold space was progressively filled by cellular matter matching the density of the subretinal debris. This transition is expected to increase resistivity of the scaffold to that of tissue, which was evidenced by concurrently improved grating acuity. These results establish a model of the controlled separation between the implant and the target neurons reproducing the features of subretinal debris in geographic atrophy of human patients, thereby providing a platform for accurate preclinical evaluation of the subretinal implants.

## Introduction

Loss of photoreceptors in diseases such as age related macular degeneration (AMD) and retinitis pigmentosa, is the leading cause of untreatable visual impairment and blindness^1^. In atrophic form of advanced AMD, called geographic atrophy (GA), the degeneration of photoreceptors in the central macula results in the loss of high-acuity vision, which severely impairs patients ability to read, recognize faces and perform other tasks of daily living^1^. While several treatment strategies aim at slowing the disease progression^2^, there are no therapies to restore vision in patients who have lost their photoreceptors. In such conditions, however, the inner retinal neurons survive to a large extent^3–5^, providing an opportunity for a functional restoration of sight by delivering visual information to these neurons via electrical stimulation. The PRIMA subretinal prosthesis (Science Corporation, California) is an array of photovoltaic pixels implanted subretinally, replacing the lost photoreceptors, to stimulate the second layer of retinal neurons – the bipolar cells. It provides central vision to patients blinded by GA with acuity of 20/420, matching its 100 μm pixel size^6–8^.

Smaller pixels are required to further improve visual acuity. However, reducing the pixel size decreases penetration depth of electric field into tissue. Therefore, performance of high-resolution implants becomes increasingly dependent on the distance between the stimulating electrodes and the target neurons. This effect is especially limiting in GA, where the target neurons in the inner nuclear layer (INL) are separated from the implant by several tens of micrometers of residual cellular debris (Figure 1A,B)^9,10^. This separation may significantly affect the stimulation threshold and spatial resolution, especially with smaller pixels.

**Figure 1.**
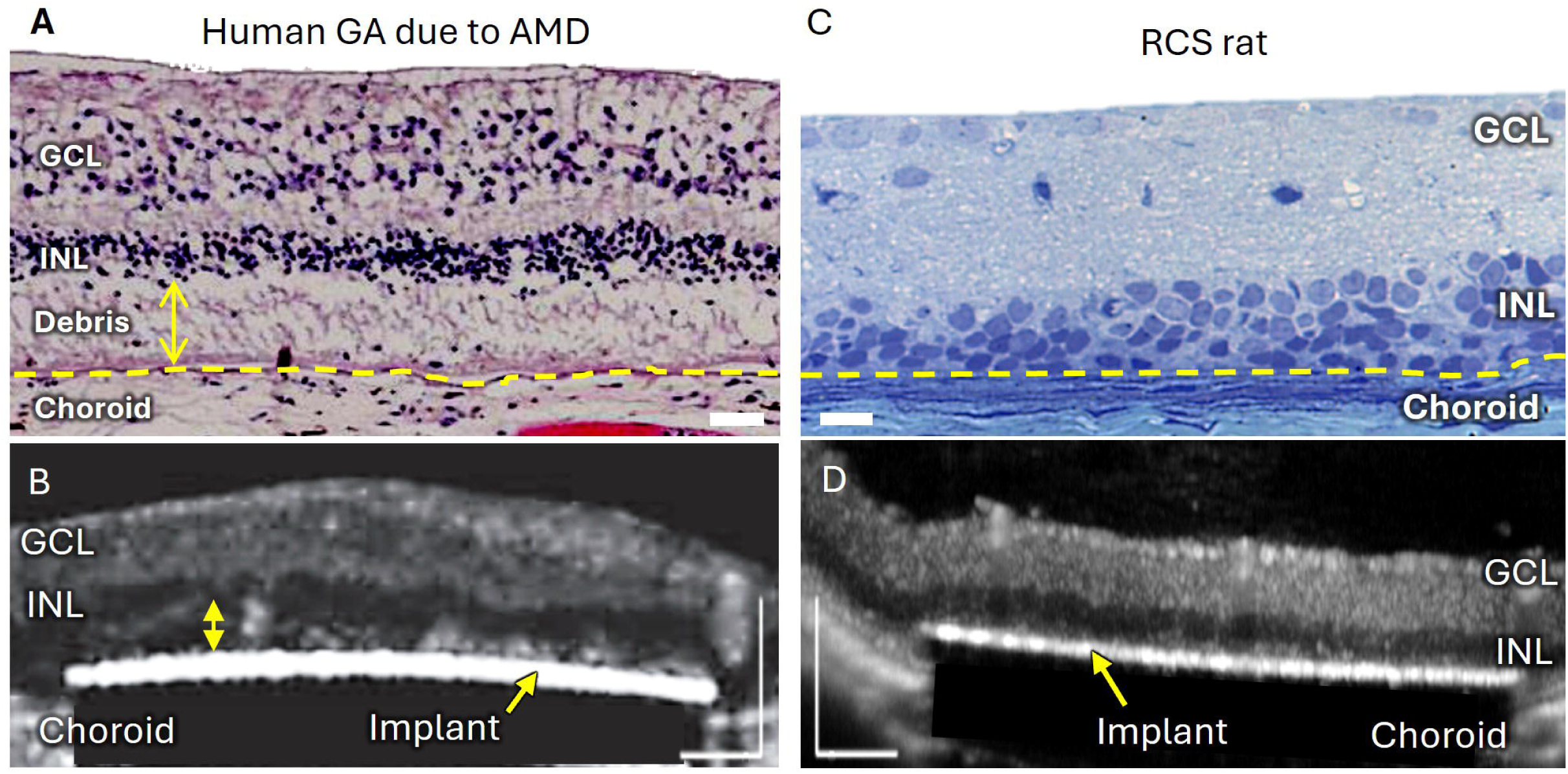
Subretinal debris in human and rodent retina. (A) Histological section of a human retina in geographic atrophy area, showing a subretinal debris layer (yellow double-headed arrow) separating the INL from the RPE. Yellow dotted lines indicate the RPE. Scale bar, 40 µm. (B) OCT image of the PRIMA prosthesis in geographic atrophy of a patient, showing persistent subretinal debris (yellow double-headed arrow) separating it from the INL. Adapted with permission from Holz *et al. N. Engl. J. Med.* (2025). Scale bar 200 µm. C) Toluidine blue-stained section of an RCS rat retina showing the inner nuclear layer (INL) directly apposed to the retinal pigment epithelium (RPE), indicated by the yellow dotted line. Scale bar, 20 µm. (D) OCT image of a PRIMA in close proximity to the INL in RCS rat, without an intervening debris layer. Scale bar 200 µm.

To address this challenge, we developed the next generation implants with smaller pixels and enhanced penetration of their electric field by current steering^11^ and by 3-dimensional electrodes ^12–15^. However, preclinical testing of the implants is impeded by absence of animal models of retinal degeneration having similar subretinal debris. In Royal College of Surgeons (RCS) rats, photoreceptor degeneration leaves the INL directly apposed to the retinal pigment epithelium and subsequently to a subretinal implant (Figure 1C,D). Retinal neurons can also migrate around three-dimensional electrodes, further altering the position of the target cells relative to the stimulating electrode^15^. Therefore, a model with controlled separation of the INL from the implant is needed for adequate preclinical evaluation of high-resolution prostheses, closely matching the clinical anatomy.

We developed a three-dimensional nanoprinted scaffold that controls the distance between the implant and the inner retinal neurons and restricts cellular migration toward the implanted electrodes. To operate on top of photovoltaic arrays, scaffolds had to meet the following requirements: they should be sufficiently transparent to allow light penetration to the implant, provide unobstructed flow of electric current, but preclude migration of cells into their 3-D structure. Moreover, to provide adequate penetration of electric field into the inner retina, resistivity of this structure should be similar to that of the retinal tissue. We determined the optimal pore size to fulfill these requirements and demonstrated that stimulation thresholds changed with the scaffold thickness similarly to that with a natural debris. This model bridges the translational gap between animal studies and the complex subretinal environment of patients with atrophic retinal disease.

## Results

### Morphological differences between rat and human degenerate retina, and their effect on electrical stimulation

In RCS rats, the inner nuclear layer (INL) is directly apposed to the retinal pigment epithelium (RPE; Fig. 1C). In contrast, in geographic atrophy of human AMD patient, a debris layer of several tens of micrometers in thickness separates the INL from RPE (yellow arrow; Fig. 1A). This difference was also evident in OCT imaging of subretinal implants: direct contact between the INL and subretinal implants in RCS rats (Fig. 1D), as opposed to a persistent debris layer separating the INL from the implant in patients with geographic atrophy (Fig. 1B).

Separation of the implant from INL strongly affects the stimulation threshold. Comparison of the voltage patterns across bipolar cell computed by RPSim simulator^16^ demonstrates how a 45 µm separation between the implant and INL, corresponding to the average debris thickness in patients, reduces the stimulus strength and spatial contrast of the pattern generated by a photovoltaic array (Fig. 2A vs B).

**Figure 2.**
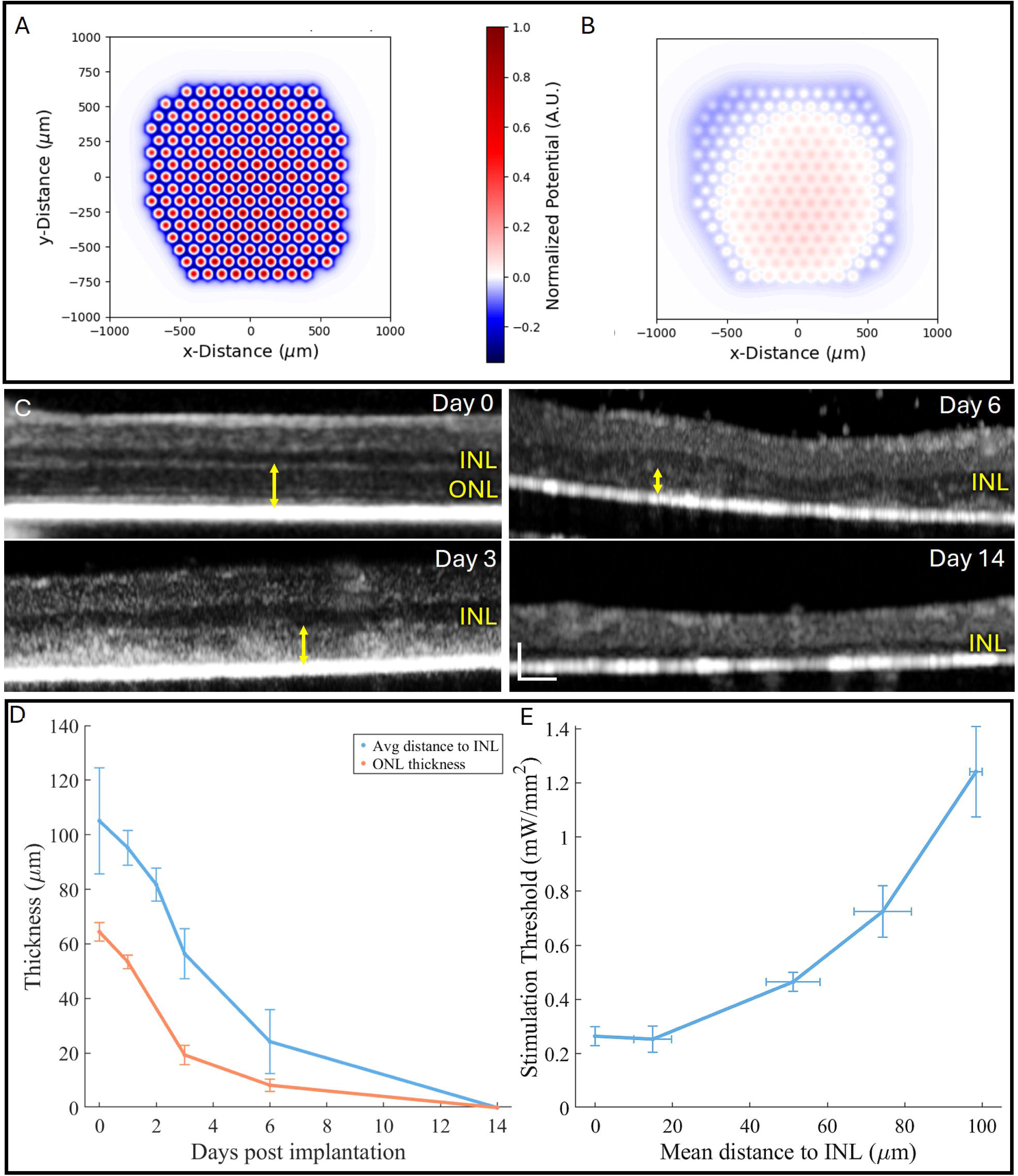
Degenerating photoreceptors as a transient model of subretinal debris. (A) Computational modeling of electric field above a photovoltaic implant at light intensity of 0.49 mW mm⁻², for retinal thickness of 170 µm and retinal tissue resistivity of 1000 Ω·cm. (B) Electric potential 45 µm above the implant, corresponding to the average thickness of subretinal debris in human geographic atrophy. (C) OCT images of Long-Evans rats demonstrating progressive degeneration of the photoreceptor layer following subretinal implantation of PRIMA devices. Scale bars 100 µm. (D) Thickness of the outer nuclear layer (ONL) thickness (n=6) and implant-to-INL distance (n = 8) extracted from the OCT images (E) Stimulation threshold as a function of the distance between the implant and INL (n=19).

Subretinal implantation into a healthy retina blocks the diffusion of oxygen and nutrients from the choroid to photoreceptors and thereby causes their degeneration above the implant. The inner retina survives since it is sustained by the inner retinal vasculature rather than choroid. Progressive degeneration of photoreceptors during several weeks post implantation in Long-Evans rats enables quantitative assessment of the effect of separation between the implant and the INL by measuring the stimulation threshold at specific timepoints. To suppress electrical stimulation of the remaining photoreceptors and ensure that responses originate from the bipolar cells, L-AP4 blocker (for ON pathway) and dark adaptation (for OFF pathway) were used during electrophysiological measurements. As shown in Figure 2C-D, OCT measurements enable quantitative assessment of progressive thinning of the outer retina between the implant and INL. At day 0, its thickness was 105±19 µm, decreasing to 56±13 µm by day 3, to 24±11 µm by day 6, and being absent by day 14, when the INL was directly apposed to the implant.

Stimulation threshold rapidly increased with larger separation between the implant and INL (Fig. 2E). Separations below 20 µm produced no change compared to a direct contact to the INL. At 45 µm, threshold increased to 0.47±0.05 mW/mm², at 75 µm, it exceeded 0.73±0.13 mW/mm², and reached 1.24±0.29 mW/mm² at 100 µm.

We attempted to replicate a subretinal debris layer in rats by performing selective photocoagulation of photoreceptors, but the resulting layer was not consistent – its thickness and its width varied significantly between laser lesions and over time (Supplementary Fig. 1A). Removal of a previously implanted device resulted in fibrotic clumps with heterogeneous and poorly controlled geometry (Supplementary Fig. 1B).

### Biomimetic scaffolds enable controlled separation between the implant and retina

To provide a layer of well-controlled thickness with resistivity matching that of the retinal tissue on top of photovoltaic arrays, we developed a porous printable scaffold. For flat implants, the scaffold was designed to maintain the INL at a defined distance above the implant surface (Fig. 3A). For pillar implants, the scaffold prevented the INL from migrating below the pillar tips, maintaining the bipolar cells at the intended stimulation range (Fig. 3B). Scaffold legs were designed to avoid contaminating the return electrodes and minimize blockage of the photosensitive area of photovoltaic pixels (Fig. 3C,D), while providing sufficient stability of the scaffold and its adhesion to the implant. Gap size of the printed mesh ranged from 3 to 7 µm to restrict migration of retinal cells (Supplementary Fig. 2A), while scaffold height provided independent control of the implant-to-retina separation (Supplementary Fig. 2B).

**Figure 3.**
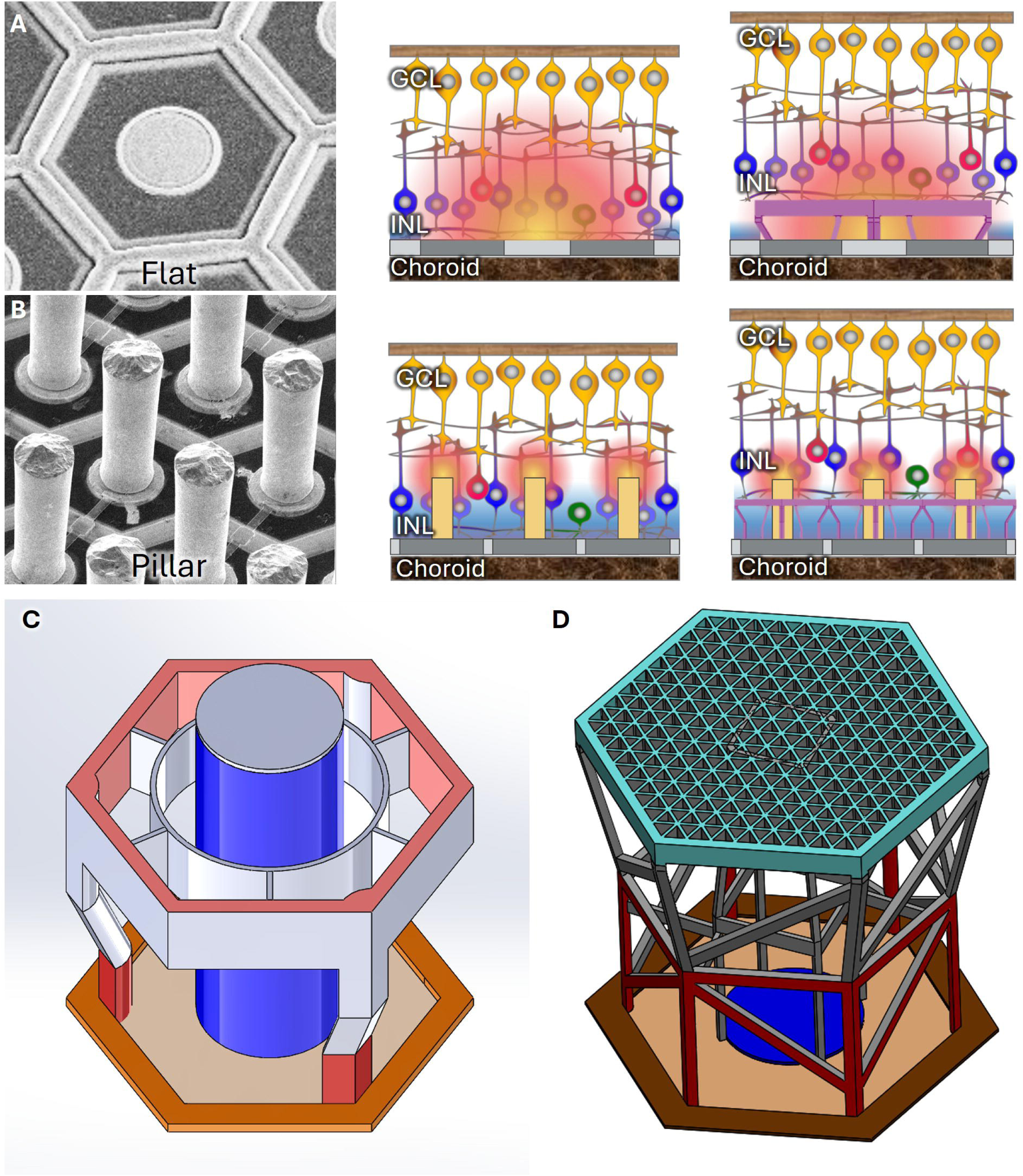
Scaffold design for flat and pillar implants. (A) SEM image of a flat photovoltaic pixel (left) and schematics illustrating the degenerate retina above the implant without (middle) and with (right) a scaffold. (B) SEM image of pillar electrodes photovoltaic implant (left) and schematics illustrating retinal integration without (middle) and with (right) a scaffold. (C) SolidWorks design of a scaffold integrated around pillar electrodes. Scaffold legs are positioned to minimize coverage of the photosensitive areas and avoid the return electrodes. (D) SolidWorks design of a scaffold above a flat photovoltaic pixel. Scaffold openings in (C) and (D) are designed to be smaller than cell bodies to restrict migration of INL through the structure.

Mechanical simulations using SolidWorks indicated necessity of cross-supports between the legs to redistribute the stress for flat structures, shown in Figure 3D and in Supplementary Fig. 3A, B. The scaffold structures were 3D printed using a two-photon polymerization technique (Methods). Laser power and scanning speed were optimized for complete and reproducible nano-printing. Low laser doses failed to create complete scaffold structures, whereas slower writing improved fabrication at the expense of print time (Supplementary Fig. 4). A laser power of 40% and writing speed of 4 mm/s provided a balance between structural fidelity and fabrication speed and were selected for subsequent fabrication. All structures were fabricated using 40x objective lens providing resolution of 1 µm and using the Nanoscribe’s IP-S photoresin with verified biocompatibility and non-cytotoxicity.

SEM imaging confirmed successful printing of scaffolds around pillar electrodes and on flat implants (Fig. 4A,B). The scaffold openings remained unobstructed at the selected print parameters. Side view with OCT further confirmed that scaffolds of 100, 45, and 15 µm in height remained stable above the implant and produced uniform separation from the device surface by the designated distance (Fig. 4C-E). It should be noted that the structures without additional reinforcements typically collapsed either in the end of the fabrication cycle or upon the implantation.

**Figure 4.**
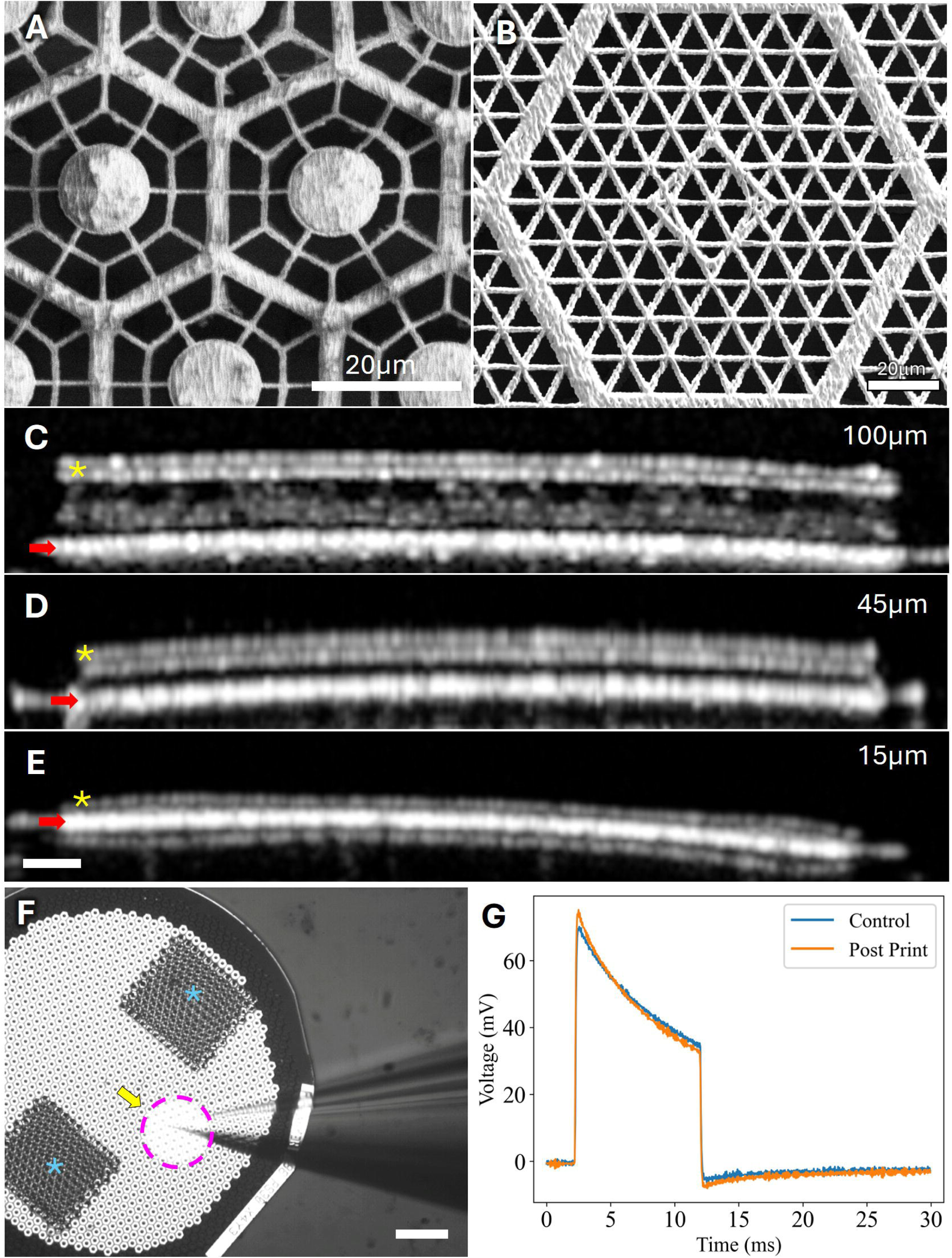
Ex-vivo assessment of the scaffolds. (A) SEM image of a scaffold printed around pillar electrodes, showing the subcellular-size gaps. Scale bar: 20 µm. (B) SEM image of a scaffold area corresponding to one pixel of a flat photovoltaic PRIMA implant. Scale bar: 20 µm. (C-E) Ex-vivo OCT images of PRIMA implants with scaffolds of 100 µm (C), 45 µm (D), and 15 µm (E) in height. Red arrows indicate the implant surface and yellow asterisks indicate the top of the scaffold. Scale bars: 100 µm. (F) Electric field mapping above the scaffold (blue asterisks) and in control regions of the same implant. Yellow arrow indicates the recording electrode and a dash circle outlines the near-infrared illumination spot. Scale bar: 200 µm. (G) Similar voltage waveforms measured above a scaffold and in a control area (n=7).

To assess the potential effect of scaffolds on optoelectronic performance of the implants in conductive medium, implants were placed in diluted saline (Hank’s balanced salt solution, HBSS) with resistivity similar to that of the retina (1,000 Ω·cm) and illuminated by pulsed NIR laser at settings similar to that of retinal stimulation: 1 mW/mm^2^, 10 ms, 2Hz. Resulting voltage waveforms were recorded at 15 µm and at 60 µm above the printed scaffolds and at similar height above intact regions of the same implant (Fig. 4F). As shown in Figure 4G, recorded waveforms were very similar between the two regions (Fig. 4G), indicating that the scaffold did not measurably impede the light delivery to the pixels, nor the current spread in electrolyte.

### Scaffold gap size controls cellular migration

To determine whether scaffolds maintained their structure under the retina and prevented retinal migration in-vivo, they were implanted subretinally in RCS rats for 6-9 weeks. Without a scaffold, the INL migrated between the pillar electrodes toward the implant surface, well below the pillar tips (Fig. 5A). A scaffold with 3 µm gaps maintained the INL, including the PKCα-positive rod bipolar cells, predominantly at and above the pillar tips, with very few nuclei observed below the scaffold (Fig. 5B). Similarly, the INL, including the bipolar cells, rested directly on flat control implants (Fig. 5C), but were maintained above the scaffold with gap sizes of 3 µm (Fig. 5D).

**Figure 5.**
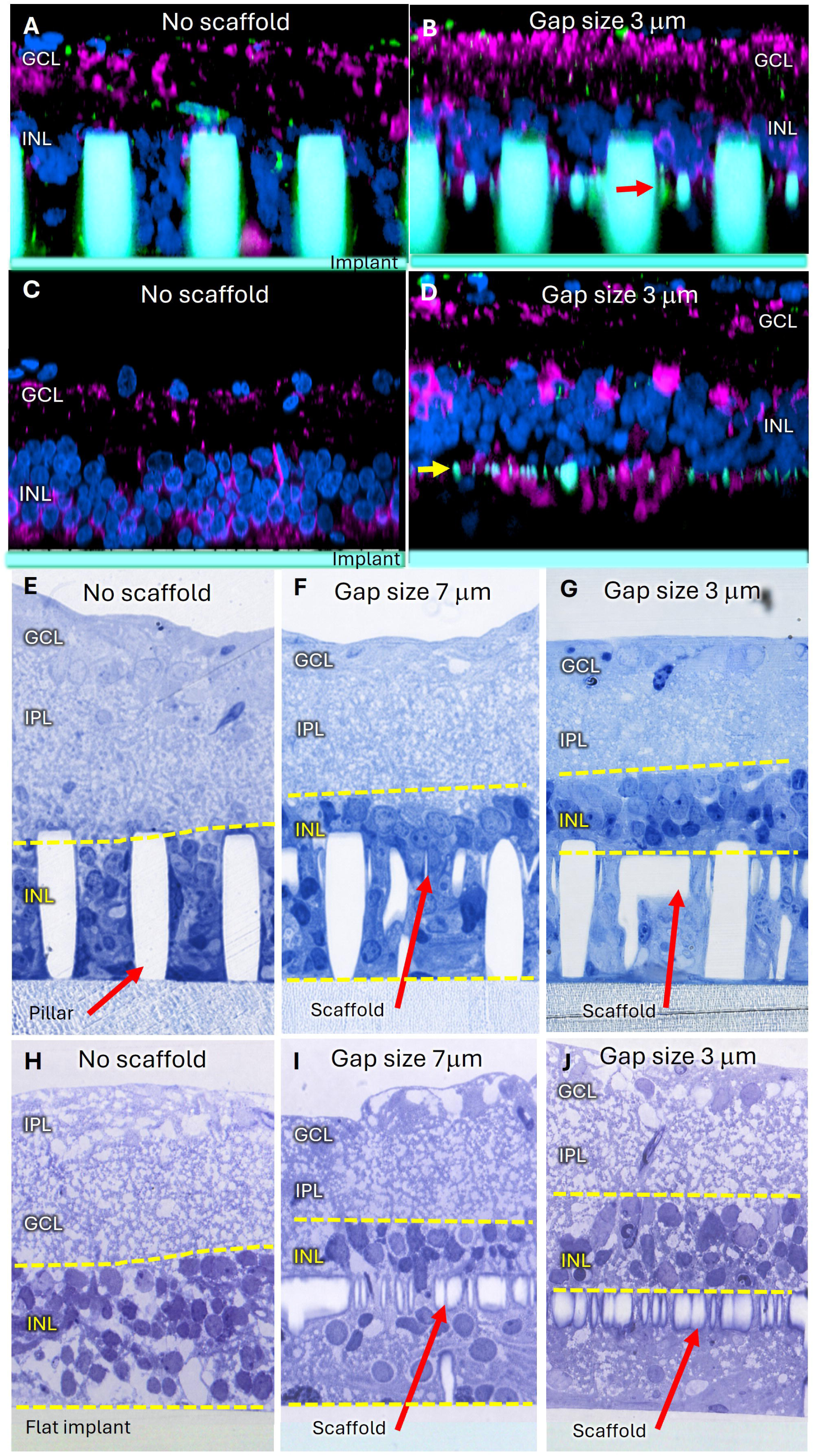
Retinal integration with scaffolds of different gap sizes. RCS rat retinas 6-9 weeks following subretinal implantation of SU-8 implants with or without printed scaffold and pillars. (A-D) confocal of immunohistochemistry where nuclei are labeled with DAPI (blue) and rod bipolar cells with PKCα (magenta). (A) Pillar implant without scaffold, showing migration of the INL between the pillars up to the implant surface (n=4). (B) Pillar implant with a scaffold (red arrow) of 3 µm gaps prevents retinal migration (n= 4). (C) Flat implant without a scaffold, where the INL, including rod bipolar cells, rest directly on the implant (n=5). (D) Scaffold with 3 µm gaps (yellow arrow) maintains retina away from the flat implant surface (n=5). (E-G) Histology of retinas implanted with pillar devices without a scaffold (E), with 7 µm scaffold gaps (F), and with 3 µm openings (G). (H-J) Histology of retinas with flat implants without a scaffold (H), with 7 µm scaffold gaps (I), or with 3 µm gaps (J). INL nuclei migrate through 7 µm gaps, but not through 3 µm.

Histology with Toluidine Blue staining confirmed retinal migration between the pillars (Fig. 5E). It also demonstrated substantial infiltration of cells below the scaffold with 7 µm gaps (Fig. 5F), but 3 µm gaps kept most of the INL above the pillar tips (Fig. 5G). The same relationship was observed with flat implants (Fig. 5H-J). Without scaffolds, INL was right at the implant surface (Fig. 5H), and many cells migrated through 7 µm openings (Fig. 5I), whereas 3 µm openings maintained the INL above the scaffold (Fig. 5J). Despite prevention of the nuclei migration, the sub-scaffold space became filled with biological material (Fig. 5G,J), which likely means that resistivity of that layer becomes similar to that of the retina above.

### Effect of scaffold on retinal stimulation

OCT imaging in rats at 6-9 weeks post implantation demonstrated that INL remained stable above the scaffolds with all heights, namely 100, 45, and 15 µm (Fig. 6A-C). VEP stimulation thresholds increased with the scaffold height (red line; Fig. 6D). With 15µm tall scaffold, there was no change relative to control implants without scaffolds, consistent with the minimal effect of small separations observed during transient photoreceptor degeneration (blue line; Fig. 6D). At larger separations, thresholds progressively increased, closely matching the threshold dependence on distance measured earlier with the transient biological debris model (blue line, Fig. 6D). The biomimetic scaffold therefore reproduced the functional effect of biological debris layer separating the INL from the implant.

**Figure 6.**
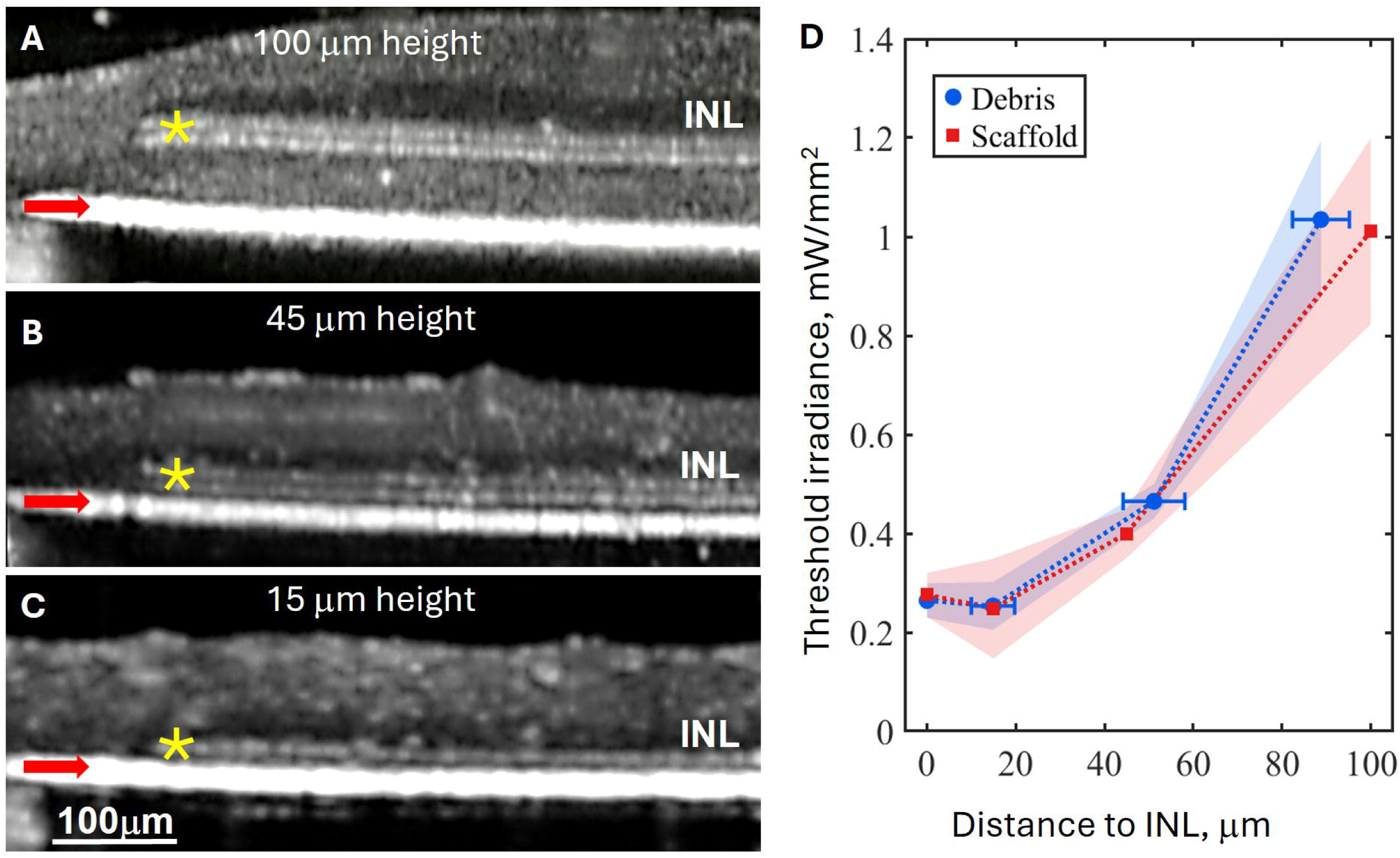
In-vivo validation of the scaffolds. (A-C) OCT images of RCS rat retinas 6-9 weeks after subretinal implantation of the PRIMA devices with scaffolds of 100 µm (A), 45 µm (B), and 15 µm (C) in height. Red arrows indicate the implants and yellow asterisks indicate the scaffolds. Scale bar: 100 µm. (D) VEP stimulation threshold as a function of distance from PRIMA implants to INL without a scaffold (0 µm; n=7) or with 15 (n=3), 45 (n=4), and 100 µm (n=4) tall scaffolds (orange). For comparison, also plotted are the thresholds measured with the transient photoreceptor degeneration model described in Figure 2 (blue).

The close agreement between the scaffold and biological debris suggested that the material occupying the sub-scaffold space reproduces the electrical properties of a natural debris layer, as has been indicated by its histological appearance (Fig. 5J). Computational modeling indicated that if the 45 µm thick sub-scaffold space would be filled with conductive physiological medium, it would shunt the electric field, attenuating the pattern amplitude and contract (Fig. 7A vs. B), which should reduce grating acuity.

**Figure 7.**
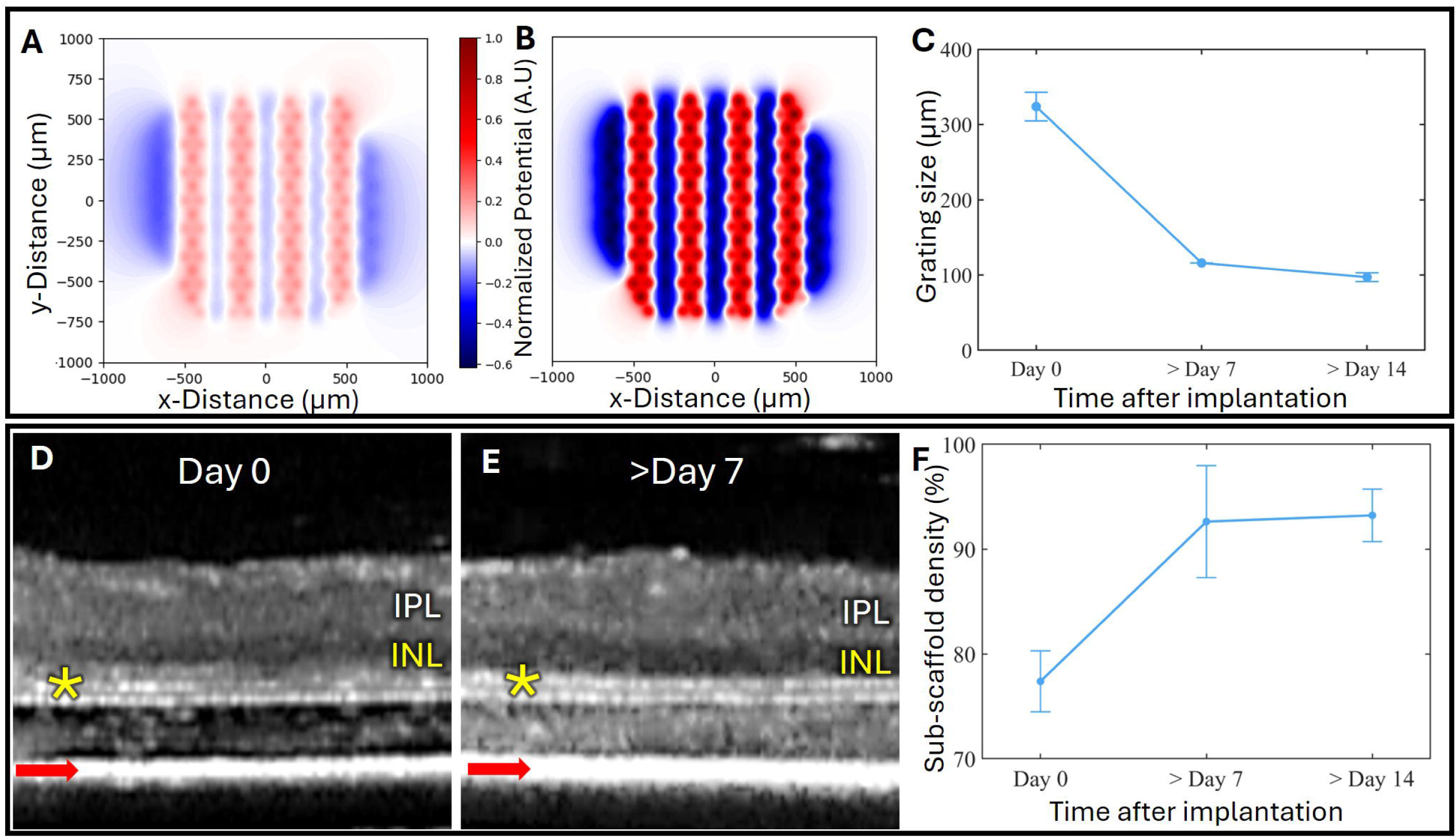
Filling the intra-scaffold space with tissue. (A-B) Simulation of electric field with the intra-scaffold space filled with 15 mS/cm saline (A) or 1 mS/cm retinal tissue (B). Simulations were performed at 3.0 mW/mm⁻² using a 154 µm grating bars, for retinal thickness of 170 µm, and a 45 µm tall scaffold. (C) Grating acuity measured with PRIMA devices having 45 µm thick scaffolds shows improvement between Days 0 and 7, which stabilizes later on (n=7). (D-E) OCT images of RCS rat retina show the intra-scaffold space immediately after implantation (D) and one week later (E). (F) OCT intensity inside the scaffold, normalized to that of the inner plexiform layer (IPL), increases from Days 0 to 7 and stabilizes later on (n=7).

To test this prediction, grating acuity was measured in RCS rats following implantation of PRIMA devices with 45 µm tall scaffolds. At the day of implantation, the threshold bar width was about 320 µm, but it improved to 100 µm at Day 14 (Fig. 7C). OCT imaging of the same retina showed a corresponding change in the sub-scaffold space: immediately after implantation, this region was hyporeflective, where only the scaffold legs scatter light, consistent with a fluid-filled space (Fig. 7D). By Day 7, OCT scattering intensity had increased in that layer, approaching that of the inner plexiform layer (IPL) (Fig. 7E). Sub-scaffold scattering intensity, normalized by that of IPL, increased from 76% at Day 0 to 93% at Day 7 and remained stable at approximately 94% at Day 14 (Fig. 7F), consistent with improvement in acuity associated with a transition from a fluid-filled space toward material with tissue-like resistivity. As seen in Figure 5D and Supplementary Figure 5, PKCα-positive rod bipolar cell processes and other biological material penetrate through the scaffold openings into the sub-scaffold region, similar to the hypertrophied outer plexiform layer in biological subretinal debris of AMD patient^10^ .

## Discussion

Age-related macular degeneration (AMD) remains the leading cause of irreversible loss of central vision in the developed world. In its advanced atrophic form, geographic atrophy (GA), degeneration of photoreceptors leaves the inner retina intact, opening the door to restoration of visual function through subretinal prostheses. The distance between the implant and the target neurons is a fundamental limiting factor of this electro-neural interface, and therefore the subretinal debris between the retinal pigment epithelium (RPE) and the inner nuclear layer (INL) in GA patients should be taken into account. However, this important feature is missing in animal models of retinal degeneration. For example, in RCS rats, the INL rests directly against the RPE and a subretinal implant. Results obtained with such preclinical models are likely to yield lower threshold and higher spatial resolution than in humans, where the residual photoreceptor material and subretinal debris separate the implant from INL by several tens of micrometers. This phenomenon is increasingly important as photovoltaic pixels are reduced in size because penetration of electric field decreases accordingly. For example, at 100 µm separation, the stimulation threshold is about 5 folds higher than in close proximity.

Degenerating photoreceptors after a subretinal implantation in WT animals can serve as a model of such a debris with variable thickness, but since this phenomenon is short-lived (only a few days), it cannot be used for reproducible longitudinal experiments. In addition, to prevent optical and electrical responses from degenerating photoreceptors, a combination of neurotransmission blockers and dark adaptation should be applied, which makes these experiments more challenging.

We addressed this challenge by developing a three-dimensional biomimetic scaffold that creates an adjustable and stable separation between the implant and the inner retina, restricts retinal migration, and reproduces the effect of biological debris layer on prosthetic vision. The scaffold’s design allows for mechanically stable separation up to 100um with no significant changes over 9 weeks of implantation. Now, the implant geometry and retinal separation can be independently controlled, enabling accurate preclinical assessment of the next generation implants.

The scaffold also resolved another very important issue with three-dimensional electrodes. Retinal neurons readily migrate into the voids of implanted structures, and in RCS rats, the INL migrated below the pillar tips, all the way to the implant surface. In humans, we expect the pillars to gradually penetrate the debris layer but remain below the INL. Therefore, uncontrolled migration of INL in RCS rats moves the target neurons into a different electric field. Restricting this migration is therefore essential in preclinical evaluation of pillar electrodes designed to remain below the bipolar cells.

Another important design requirement was that the scaffold should not compromise penetration of light to the implant and spread of electric field from the electrodes. Selection of a transparent material and the porous architecture addressed both requirements, as demonstrated by similar waveforms generated by the implants with and without a scaffold ex-vivo. Scaffold also closely reproduced the effect of biological debris on stimulation thresholds in-vivo. Biological material filling the scaffold after implantation appears similar to the inner plexiform layer, which contributes to accuracy of this preclinical model in replicating a subretinal biological structure in AMD patients. Since we could not measure resistivity of this layer in-vivo directly, we can only suggest that similarity based on the matching electrophysiological measurements. It is also important to keep in mind that human subretinal debris is heterogeneous and include basement membrane deposits, CD68-positive macrophages, localized subretinal fibrosis, and disease-associated extracellular components^10^ that may not be represented inside the scaffold.

The utility of this model extends beyond the photovoltaic arrays - biomimetic scaffolds can be applied to any flat or 3-dimensional subretinal implants to ensure proper separation from the target neurons while preventing the migration-induced loss of neural structure, which is essential for representative preclinical testing.

## Methods

### Human retinal histology

Previously anonymized human retinal sections stained with hematoxylin and eosin (H&E) were obtained from the Stanford teaching slide library. Sections from eyes with age-related macular degeneration (AMD) were imaged by light microscopy to assess the presence of subretinal debris separating the inner retina from the retinal pigment epithelium.

### Biological models of persistent subretinal separation

Two biological approaches were evaluated to generate a persistent tissue layer separating the inner retina from the retinal pigment epithelium (RPE). First, subretinal fibrosis was induced in Long-Evans rats using a two-stage laser photocoagulation protocol adapted from previously described models of subretinal fibrosis^17^. Laser treatment was performed using a PASCAL pattern scanning laser photocoagulator (PASCAL; Topcon Medical Laser Systems, Santa Clara, CA, USA). Laser lesions were generated using a 532 nm wavelength, 100 µm spot diameter, 100 ms pulse duration, and 250 mW power. A second laser exposure was applied to the same lesions 7 days later using the same parameters to promote a persistent fibrotic response. The resulting subretinal lesions were monitored longitudinally by OCT to assess their thickness, spatial distribution, and persistence.

The second approach was based on the tissue response following subretinal implantation and subsequent extraction in RCS rats^14^. Six weeks after subretinal implantation, devices were surgically removed and the retina was allowed to recover for another six weeks. OCT was used to monitor the formation and persistence of tissue within the subretinal space following the implant extraction.

Both approaches generated persistent subretinal material but with substantial variability in its thickness and spatial distribution, which precluded their application in studies where separation between the implant and the INL should be carefully controlled.

### Simulations of electric field

Electric field was modeled using the Retinal Prosthesis Simulator (RPSim), an in-house simulation framework previously developed for photovoltaic retinal prostheses^16^. RPSim combines finite element modeling of the implant-generated electric field in COMSOL Multiphysics with a SPICE-based circuit solver implemented in Xyce to account for the nonlinear electrical behavior of individual photovoltaic pixels coupled through the common conductive medium. Implant geometry and electrical parameters were based on the previously validated PRIMA model^16^.

To evaluate the effect of separation on stimulation, electric potential generated by a PRIMA implant was simulated with the retina positioned directly above the implant and separated by a 45 µm layer, corresponding to the average thickness of debris observed clinically. Simulations used an incident irradiance of 0.49 mW mm⁻² and a retinal thickness of 170 µm. The layer was assigned the same electrical conductivity as that of retinal tissue: 1 mS cm⁻¹. Electric potential distribution was calculated immediately above the implant and at the retinal interface to determine the effect of separation on stimulus strength and spatial contrast.

A second set of simulations was performed to assess the effect of conductivity of the sub-scaffold space on stimulation. A 45 µm thick scaffold was modeled beneath a 170 µm thick retina using a 154 µm alternating grating pattern at an irradiance of 3.0 mW mm⁻². The sub-scaffold compartment was assigned either the conductivity of saline (15 mS cm⁻¹) or of the retina (1 mS cm⁻¹). RPSim implementation and validation have been described previously^16^.

### Implants fabrication

PRIMA implants with 100 µm photovoltaic pixels and local return electrodes surrounding each pixel were used in this study. The 1.5 mm diameter implants were fabricated by Pixium Vision as previously described^6^. The devices are 30 µm thick and optimized for near-infrared illumination at 880 nm. Each photovoltaic pixel converts incident light into electrical current delivered through an active electrode and a surrounding local return electrode.

Three-dimensional pillar electrodes were fabricated on 4-inch p-doped silicon wafers by electrodeposition of gold using a gold plating solution (NB Semiplate AU 100TH; NB Technologies, Bremen, Germany), as described previously^15^. Photolithographically defined patterns for both active and return electrodes were first created using a thin adhesion layer of titanium (50 nm) followed by a gold seed layer (200 nm) deposited via sputtering (Kurt J. Lesker, USA). These interconnections permitted uniform electroplating across the entire wafer. Electrodes were formed into pillar-shaped structures by applying a constant current density of 1 mA/cm², yielding a deposition rate of approximately 3 µm/h. For the 40 µm-wide pixels used in this study, this process produced gold pillars with a diameter of 16.5 µm and the desired height. Following electroplating, the exposed electrode surfaces were coated with a bilayer of titanium (40 nm) and sputtered iridium oxide (SIROF; 436 nm) to increase charge injection capacity (EIC Laboratories, MA, USA). For in vivo anatomical studies, a 200 nm titanium coating was deposited over the entire implant to enhance mechanical stability and biocompatibility prior to subretinal implantation^15^.

### Animals

All procedures involving animals were approved by the Stanford Administrative Panel on Laboratory Animal Care and conducted in accordance with the ARVO Statement for the Use of Animals in Ophthalmic and Vision Research. Royal College of Surgeons (RCS) rats from a breeding colony maintained at Stanford University were used. Animals were older than postnatal day 180 at the time of implantation to ensure complete photoreceptor degeneration.

Rats were anesthetized with intraperitoneal ketamine (75 mg/kg) and xylazine (5 mg/kg). After pupil dilation and preparation of the surgical field, a 1.5 mm scleral incision was made approximately 1 mm posterior to the limbus. Sterile saline was injected into the subretinal space to create a localized retinal detachment. The implant was introduced through the incision and positioned at least 3 mm from the retinotomy. The conjunctiva was closed with 10-0 nylon sutures, followed by application of topical antibiotic ointment.

For histological studies, implants remained in the subretinal space for six weeks before tissue collection. Animals were euthanized by intracardiac administration of pentobarbital/phenytoin solution. Eyes were immediately enucleated and rinsed in phosphate-buffered saline (PBS). The cornea, iris, and lens were removed, and the posterior eyecup containing the implant was isolated. For whole-mount immunohistochemistry, the retinal region surrounding the implant was dissected and fixed overnight in 4% paraformaldehyde at 4 °C.

### Scaffold printing

Scaffold geometries were designed in SolidWorks (Dassault Systèmes) and fabricated by two-photon lithography (Nanoscribe GmbH & Co. KG, Karlsruhe, Germany) directly onto SU-8 polymer discs and monopolar and PRIMA photovoltaic arrays.

To evaluate the mechanical stability of the designed structures, finite element analysis was performed in SolidWorks Simulation (Dassault Systèmes) to assess von Mises stress distributions and buckling behavior for an individual 100 μm pixel. Models were fixed at the interfaces between the scaffold legs and the underlying surface, and material properties were assigned based on manufacturer specifications for Nanoscribe IP-S photoresist (Nanoscribe GmbH & Co. KG, Karlsruhe, Germany). A 2 kPa pressure was applied vertically to the scaffold surface, corresponding to a total force of approximately 7 μN. To model asymmetric forces within the subretinal space, lateral forces of either 7 μN or 1 mN were applied concurrently with the vertical pressure. Both the original and reinforced scaffold geometries were evaluated under each lateral loading condition. Von Mises stress distributions and buckling analyses were used to assess mechanical stress and structural stability under each loading condition (Supplementary Figure 3).

Nanoscribe fabrication parameters were optimized using a laser dosage grid process in which laser writing speed and laser power were systematically varied. Writing speeds of 2, 4, 8, 12, and 16 mm/s were tested at laser powers of 20%, 30%, 40%, 50%, and 60%. For each parameter combination, complete scaffold cells were fabricated to assess the overall print fidelity, including resolution of the upper structural features. In parallel, the scaffold legs were printed independently onto the substrate to evaluate fabrication quality. Samples were examined by light microscopy and Scanning Electron Microscopy (SEM) to identify parameter combinations that produced complete, well-defined scaffold geometries while maintaining consistent contact between the scaffold legs and the underlying surface (Supplementary Figure 4). These results were used to select the laser power and writing speed for subsequent scaffold fabrication.

All the scaffold fabrication was performed using 40x objective with resolution of 1µm. Before printing on photovoltaic arrays, implants were cleaned in acetone and isopropyl alcohol, then secured on a PDMS-coated wafer to maintain position during fabrication processes. A drop of IP-S photoresin (Nanoscribe GmbH & Co. KG, Karlsruhe, Germany) was applied to the printing surface, and the wafer was loaded into the Nanoscribe system. IP-S photoresin is a general-purpose, acrylate-based negative-tone photoresin with verified biocompatibility and non-cytotoxicity as confirmed by the manufacturer. After printing, samples were developed in SU-8 Developer for 2 h followed by a 15 min isopropyl alcohol rinse, with agitation during the SU-8 Developer step to facilitate removal of excess resin from narrow gaps. Prints were finally treated with oxygen plasma (plasma ashing, 30s at 15sccm O₂, 150 W; SamCo) to increase surface hydrophilicity of the polymerized IP-S structures before subsequent implantation.

### Transparency Test

To assess whether the printed scaffold altered light transmission or electric current propagation, photovoltaic implants containing patches of 45 µm-high scaffolds were immersed in diluted saline solution with a resistivity of approximately 1,000 Ω·cm. NIR spots (1 mW/mm², 10 ms pulses, 2 Hz) were projected onto scaffold and control regions, and the resulting electric potential was measured using an electrode positioned 47 µm above the implant surface. For each implant, voltage amplitudes measured over scaffold spots were normalized to the voltage amplitude measured over the unprinted control region of the same implant. These normalized values were then used for statistical significance testing to determine whether scaffold-covered regions differed from control.

### Optical coherence tomography

To assess the success and integrity of the print, spectral-domain optical coherence tomography (OCT; HRA2-Spectralis, Heidelberg Engineering, Heidelberg, Germany) was used to obtain cross-sectional views of the printed scaffolds on top of the implants. Implants glued to PDMS layers were positioned vertically in front of the OCT module and imaged in air.

To monitor the implant-retina interface and retinal morphology, OCT was performed following implantation and longitudinally during the experimental period. The implant position, retinal reattachment, and morphology were assessed during 6 weeks post implantation. For imaging, the cornea was covered with viscoelastic gel and a coverslip was positioned over the eye to compensate for corneal curvature and refractive power.

To monitor the degeneration of photoreceptors in the transient debris model, OCT images were taken daily to determine the degeneration timeline. Since degeneration is a non-uniform event, measurements were taken as an average across 25 points above the implant.

### Immunohistochemistry and confocal imaging

Fixed retinal samples were permeabilized for 3 h at room temperature in PBS containing 1% Triton X-100 and subsequently blocked with 10% bovine serum albumin (BSA). Samples were incubated for 48 h at room temperature with mouse anti-PKCα antibody (Santa Cruz Biotechnology, sc-8393) diluted in PBS containing 5% BSA and 0.5% Triton X-100. After washing with PBS containing 0.1% Tween-20, samples were incubated for 48 h at room temperature with Cy3-conjugated donkey anti-mouse secondary antibody (Jackson ImmunoResearch, 715-165-150). Nuclei were counterstained with DAPI before final washing and mounting in antifade medium.

Retinal whole mounts were imaged using a Zeiss LSM 880 laser-scanning confocal microscope (Carl Zeiss, Germany). Reflected light was used to visualize the implant surface. Z-stacks spanning the retinal thickness from the inner limiting membrane to below the implant surface were acquired using a 40x oil-immersion objective. Images were processed using Fiji/ImageJ. Acquisition parameters were maintained across experimental groups for each staining condition.

### Scanning electron microscopy

Implants and printed scaffold structures were characterized using scanning electron microscopy (Zeiss Sigma SEM with Schottky Field Emission (FE) source and GEMINI electron optical column; Zeiss, Oberkochen, Germany). Samples were mounted on SEM holders using conductive carbon tape and imaged under vacuum. Low accelerating voltages were used to visualize the implant and scaffold surfaces while minimizing electron penetration into the underlying structures. SEM imaging was used to assess scaffold geometry, structural integrity, pore dimensions, clogging issues and integration of the printed structures with flat and pillar implants.

### Histological preparation

Eyes collected for histology were fixed in 1.25% glutaraldehyde for 24 h at room temperature and post-fixed in osmium tetroxide for 2 h. Samples were dehydrated through graded alcohol and propylene oxide and infiltrated overnight at room temperature with Araldite-EMbed epoxy resin without DMP-30 (RT13940, Mollenhauer’s kit; Electron Microscopy Sciences). Samples were subsequently embedded and cured at 70 °C for 36 h.

Epoxy blocks were trimmed to expose the implanted retinal region. Sections were cut at 700 nm using a Reichert Ultracut E ultramicrotome and stained with 0.5% toluidine blue for light microscopy.

### Electrophysiology

Visually evoked potentials (VEPs) were recorded using transcranial electrodes implanted above the visual cortex. One electrode was positioned over each hemisphere of the primary visual cortex (V1; 4 mm lateral to the midline and 6 mm caudal to bregma), with a reference electrode positioned over the somatosensory cortex (2 mm right of the midline and 2 mm anterior to bregma).

Animals were lightly anesthetized and pupils were dilated before recording. Artificial tear gel and a coverslip were placed over the cornea to compensate for its optical power. Optical patterns were projected onto the subretinal PRIMA implant using a customized projection system consisting of an 880 nm near-infrared (NIR) laser (MF_880 nm_400 um; DILAS, Tucson, AZ), collimating optics, a digital micromirror display (DMD; DLP Light Commander, LOGIC PD, Carlsbad, CA), a slit lamp (SL-120; Carl Zeiss, Thornwood, NY), and a NIR-sensitive CCD camera (acA1300-60gmNIR; Basler, Ahrensburg, Germany). Retinal irradiance was calculated from the ratio of the projected pattern dimensions at the cornea and retina.

For stimulation threshold measurements, NIR pulses of 10 ms duration were delivered at 2 Hz over a range of irradiances. VEPs were recorded using an Espion E3 system (Diagnosys LLC, Lowell, MA) at a sampling rate of 2 kHz and averaged over 250 trials. Peak-to-peak VEP amplitude was calculated as a difference between the first trough in the signal within 120 ms of the stimulus onset and the peak within 100 ms after the trough. Noise was determined in a similar fashion, 320 ms after the stimulus onset. The stimulation threshold was defined as the irradiance required to obtain a signal-to-noise ratio (SNR) of 1 through linear extrapolation of two points with an SNR > 1. The threshold was checked by ensuring that at irradiance below the extrapolated threshold, no signal was detected (SNR < 1)^18^. Implant-to-INL distance was measured by OCT and used to determine the relationship between retinal separation and stimulation threshold. Measurements were performed during transient photoreceptor degeneration and after implantation of PRIMA devices with scaffolds of different heights.

For measurements in Long-Evans rats during transient photoreceptor degeneration, pharmacological blockade and dark adaptation were used to isolate prosthetic responses from those generated via residual photoreceptors by light or direct electrical stimulation. L-AP4 was administered before VEP recording by intravitreal injection to block ON pathway, and measurements were performed under dark-adapted conditions to prevent electrical response of photoreceptors via OFF pathway. This allowed the remaining outer retina to function as inert separation layer while minimizing visually evoked responses mediated by native photoreceptors.

Grating acuity was measured by projecting alternating grating patterns onto PRIMA implants. NIR stimulation was delivered at 4 mW mm⁻² using 4 ms pulses at a carrier frequency of 64 Hz. Grating polarity was alternated at 2 Hz, corresponding to 500 ms per pattern. Grating bar width was progressively reduced to determine the smallest pattern that elicited a VEP response. Grating threshold was determined in a similar fashion as mentioned above.

VEP data were analyzed using custom MATLAB scripts. High-frequency components associated with the 64 Hz carrier were removed using a spectrum reconstruction algorithm. Grating acuity was determined from the intersection between a logarithmic fit to responses above an SNR of 1.

### Quantification of cellular migration

To assess retinal migration through scaffolds of different gap sizes, SU-8 polymer discs containing 300 × 300 µm scaffold regions with 45 µm height and gap sizes of 9, 7, and 5μm were implanted subretinally. Following enucleation, eyes were processed for histology, sectioned through the implanted regions, and stained with toluidine blue. Sections containing the scaffold were imaged, and INL cells located above and below the scaffold surface mesh were counted. For each section, the ratio of INL cells above and below the scaffold was calculated and compared across scaffold gap sizes to quantify percent of INL migration.

To quantify changes in the optical properties of the sub-scaffold space over time, OCT images were acquired at multiple timepoints following implantation of devices with scaffolds of 100μm height. Raw OCT images were analyzed in ImageJ using fixed-size rectangular regions applied consistently across samples. Sub-scaffold intensity was measured within the region directly beneath the scaffold and above the implant surface using the ImageJ histogram tool. The same-size reference region was placed within the inner plexiform layer (IPL) above the scaffold, avoiding contact with the scaffold and INL. Mean sub-scaffold intensity was normalized to the corresponding IPL intensity from the same OCT image and expressed as a percentage of IPL intensity for comparison across timepoints.

### Data and code availability

All relevant data supporting the key findings of this study are available within the article and its Supplementary Information. Any additional data that support the findings of this study are available from the corresponding author upon reasonable request.

Codes were previously published in https://doi.org/10.1038/s41467-022-34353-y and are available from the corresponding author upon reasonable request.

## Competing interests

M.B.B, D.P. and T.K. serve as consultants to Science Corp (formerly Pixium Vision). D.P.’s patents related to retinal prostheses are owned by Stanford University and licensed to Science Corp, the details of which are disclosed below. All other authors declare no competing interests. Patent: - patent applicant: Stanford University - name of inventor(s): D. Palanker, A. Vankov, M. Blumenkranz - application number: US 7,047,080 - status of application: issued - specific aspect of manuscript covered in patent application: use of photovoltaic pixels for retinal prosthetics.

## Acknowledgement and contributions

DP and MB conceived and supervised the project, MB performed surgeries; MB, VM, DPPH performed in-vivo imaging, immunohistochemistry and confocal imaging; DPH, VM and MB conducted image processing and statistical analysis; VM, VS and LG fabricated subretinal implants and printed scaffolds under the guidance of TK; RD sectioned the retina; MB and DP guided the research and data analysis; MB, VM and DPH assembled the figures; MB and DP wrote the manuscript, all other authors reviewed the manuscript. Studies were supported by BrightFocus Foundation (Grant M2025007N), the NIH (Grants R01-EY-035227, P30-EY-026877), the Department of Defense (Grant W81XWH-2210933), and AFOSR (Grant FA9550-19-1-0402, FA9550-24-1-0138), and unrestricted grant from the Research to Prevent Blindness, Wu Tsai Institute of Neurosciences at Stanford. Photovoltaic arrays were fabricated at the Stanford Nano Shared Facilities (SNSF) and Stanford Nanofabrication Facility (SNF), which are supported by the National Science Foundation award ECCS-2026822.

## Notes

### Competing Interest Statement

M.B.B, D.P. and T.K. serve as consultants to Science Corp (formerly Pixium Vision). D.P. patents related to retinal prostheses are owned by Stanford University and licensed to Science Corp

