## Supplemental Figures for "Biomimetic scaffolds mimic human subretinal debris for preclinical testing of prosthetic vision"

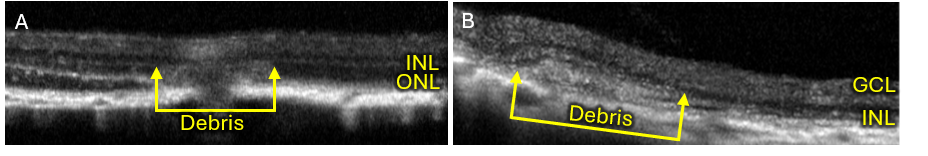

**Supplementary Figure 1. Biological models of subretinal debris.** (A) OCT image of a Long-Evans rat following double laser photocoagulation to induce subretinal fibrosis. Yellow arrows indicate the fibrotic layer formed between the retina and RPE. Scale bar, 500 µm. (B) OCT image of an RCS rat following explantation of a subretinal implant. Yellow arrows indicate the subretinal fibrotic layer formed after the device removal. Scale bar, 500 µm.

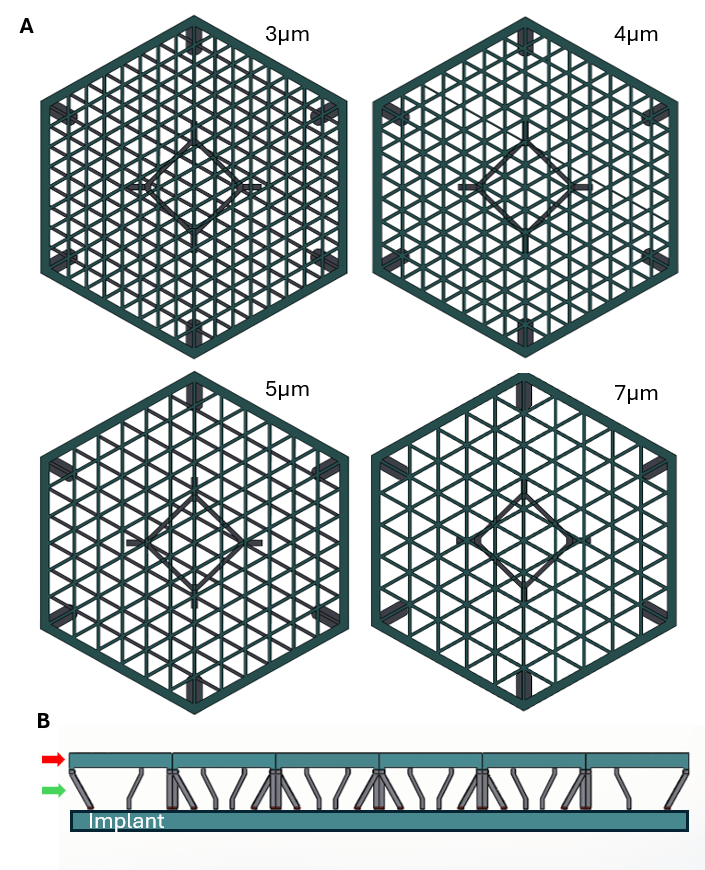

**Supplementary Figure 2. Scaffold geometry and dimensions.** (A) Top-down view of the scaffold design over a 100 µm flat photovoltaic pixel showing scaffold openings of 3, 4, 5, and 7 µm used to evaluate the effect of gap size on retinal cell migration. (B) Side view of the scaffold design showing its position relative to the underlying implant. The red arrow indicates the top of the scaffold and the green arrow indicates the scaffold legs.

**Supplementary Figure 3. Mechanical simulation of scaffold leg designs.** SolidWorks Simulation was used to compare von Mises stress distributions and buckling stability in the original scaffold geometry with separated legs (A) and a reinforced design incorporating additional cross-supports (B). Modeling was performed for a single 100 μm pixel, with fixed points defined at the interfaces between the scaffold legs and the underlying surface and material properties assigned according to manufacturer specifications for Nanoscribe IP-S photoresist. A 7 μN force (2 kPa pressure) was applied vertically to the scaffold surface. Under vertical loading alone, the unbraced structure experienced greater mechanical stress and had approximately half the factor of safety of the reinforced design. A lateral force was then applied to represent asymmetric loading in the subretinal space. An equal-magnitude 7 μN lateral force induced buckling of the unbraced structure, whereas the reinforced design remained stable even under a substantially larger 1 mN lateral force, as shown in B.

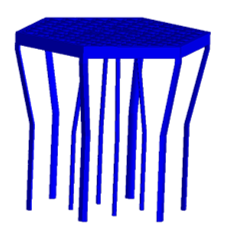

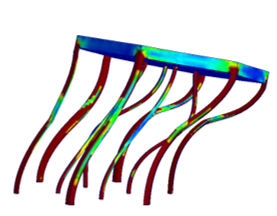

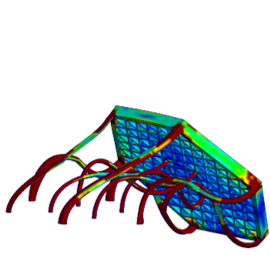

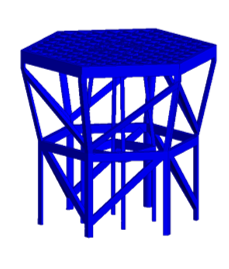

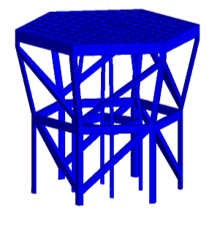

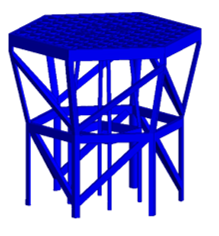

**A**

**B**

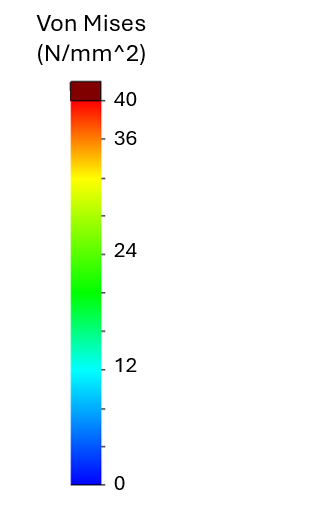

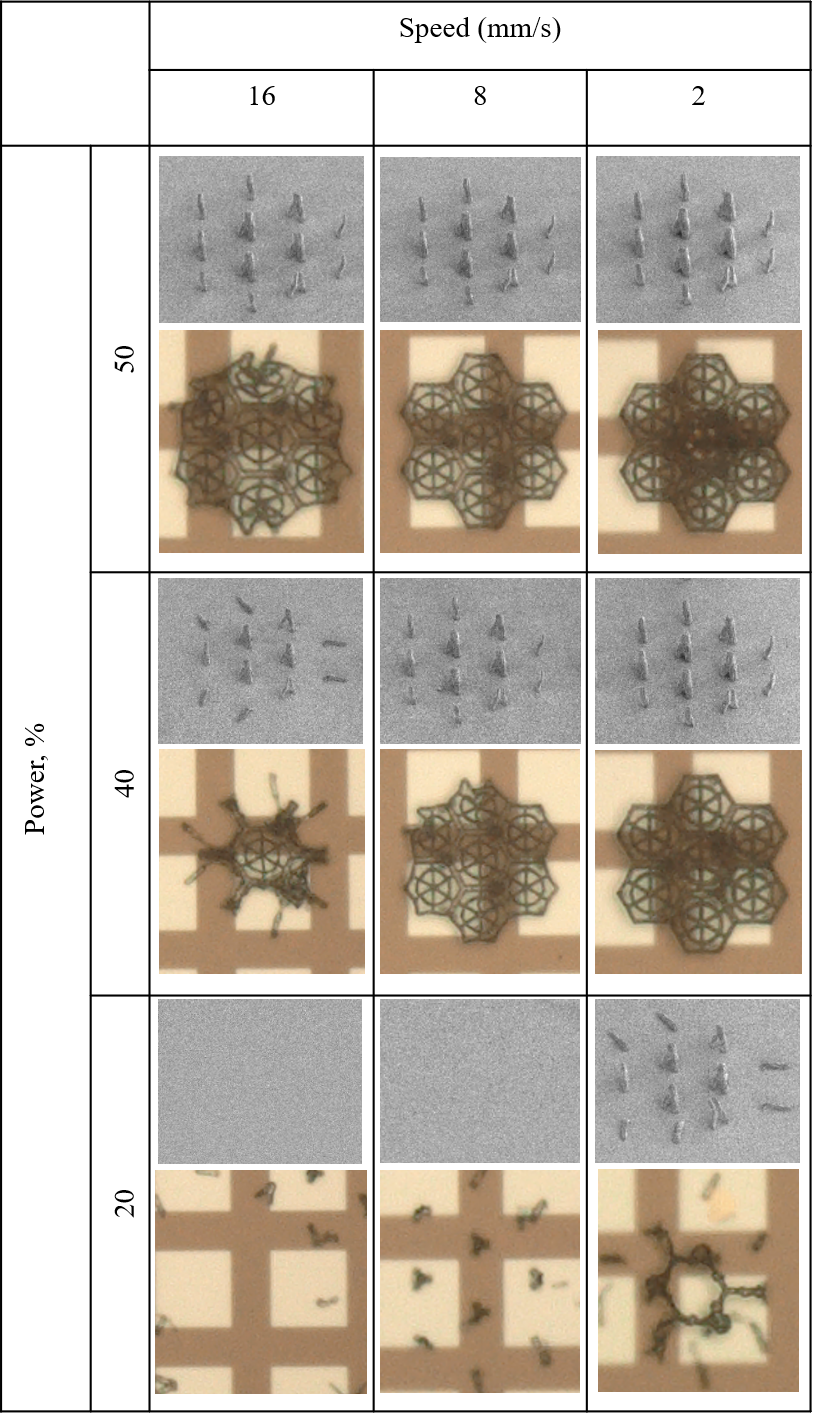

**Supplementary Figure 4. Laser dosage optimization for two-photon polymerization of scaffold structures.** Representative laser dosage matrix showing scaffold fabrication across selected combinations of laser power and printing speed. Complete scaffold structures and individual scaffold legs were printed at each set of parameters to assess structural resolution, print fidelity, and attachment to the underlying substrate. Images show representative structures obtained across the laser dose range.

**Supplementary Figure 5. Sub-scaffold material penetrates through the scaffold mesh.** (A-B) Toluidine blue-stained sections following implantation of scaffolds with porous (A) or solid (B) sidewalls show similar sub-scaffold filling. (C) Immunofluorescence image of a scaffold-implanted retina showing rod bipolar cells labeled with PKCα and the scaffold indicated by the red arrow. Magenta processes corresponding to bipolar cells dendrites extend into the sub-scaffold space.

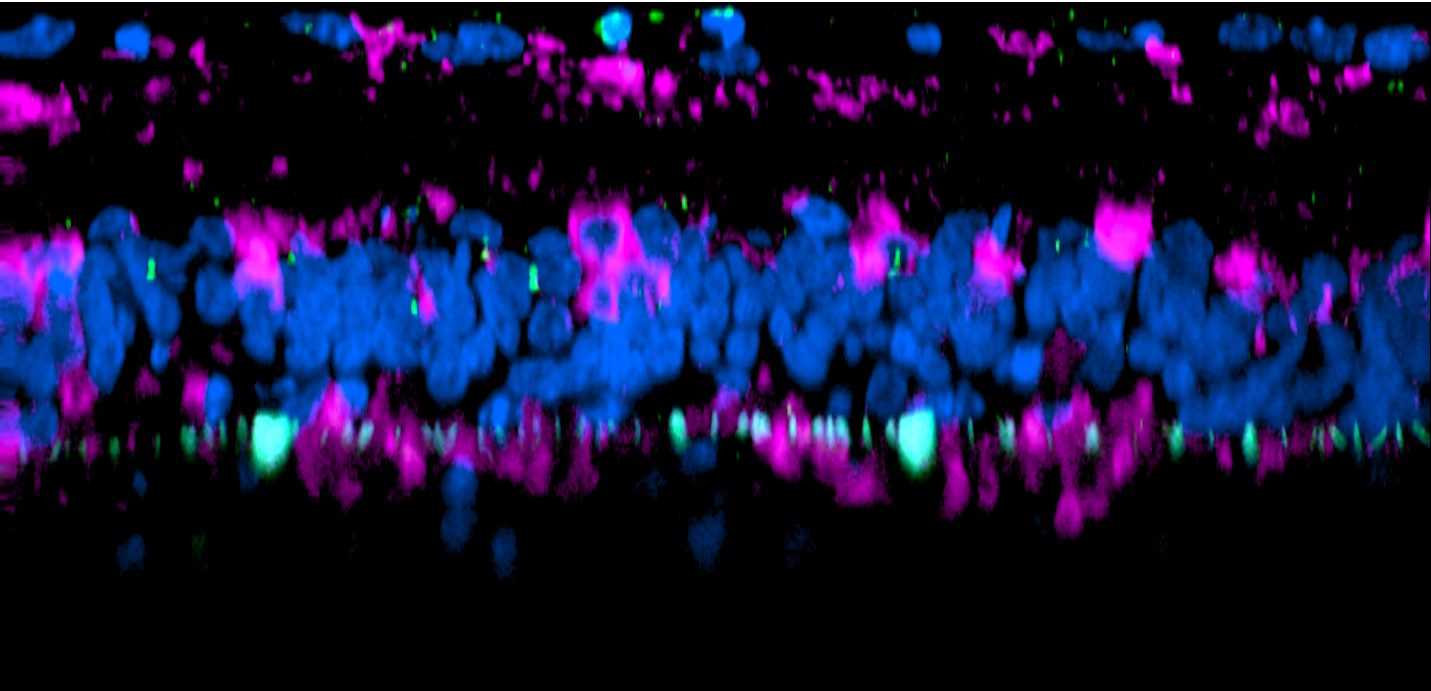

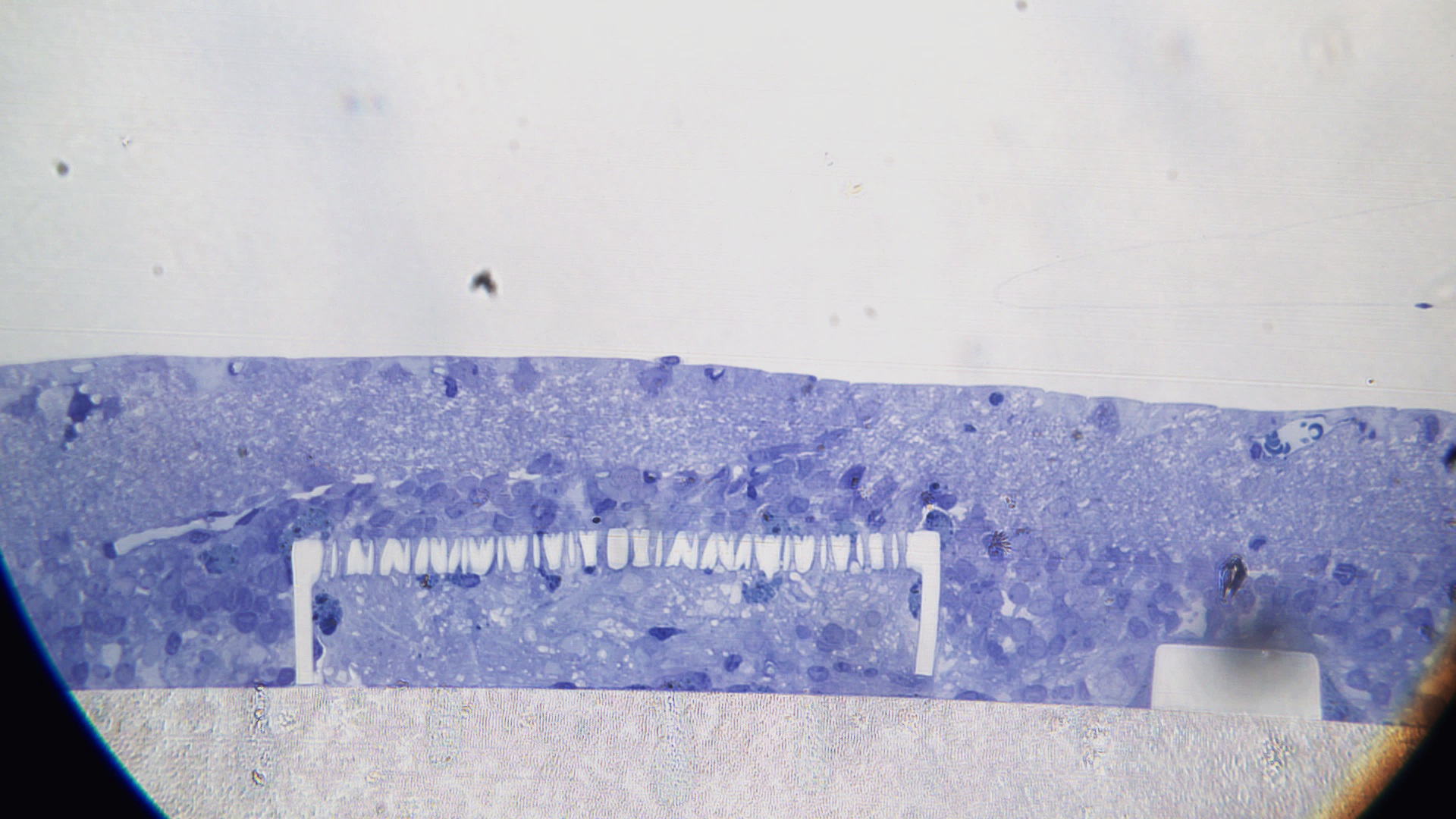

Solid side wall

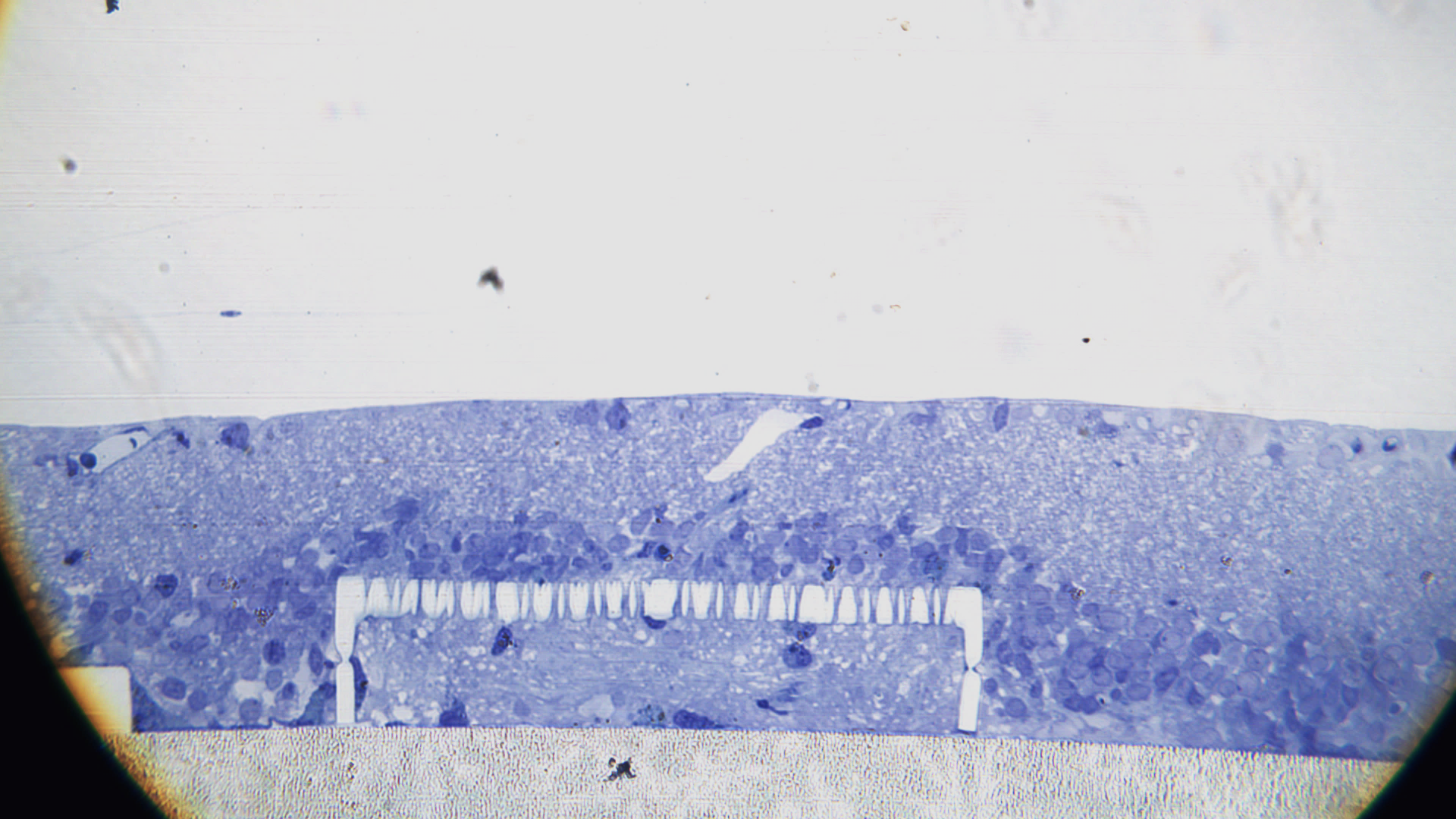

Porous side wall

GCL

IPL

INL

Sub-scaffold tissue

GCL

IPL

INL

Sub-scaffold tissue

**A**

**C**

GCL

INL

**B**
